# Human Intelligence and the Connectome are Driven by Structural Brain Network Control

**DOI:** 10.1101/2023.08.02.551642

**Authors:** Evan D. Anderson, Lav R. Varshney, Babak Hemmatian, Pablo D. Robles-Granda, Anuj Keshava Nayak, Ramsey R. Wilcox, Christopher E. Zwilling, Been Kim, Aron K. Barbey

## Abstract

Research in network neuroscience demonstrates that human intelligence is shaped by the structural brain connectome, which enables a globally coordinated and dynamic architecture for general intelligence. Building on this perspective, the network neuroscience theory proposes that intelligence arises from system-wide network dynamics and the capacity to flexibly transition between network states. According to this view, network flexibility is made possible by network controllers that move the system into specific network states, enabling solutions to familiar problems by accessing nearby, easy-to-reach network states and adapting to novel situations by engaging distant, difficult-to-reach network states. Although this framework predicts that general intelligence depends on network controllability, the specific cortical regions that serve as network controllers and the nature of their control operations remain to be established. We therefore conducted a comprehensive investigation of the relationship between regional measures of network controllability and general intelligence within a sample of 275 healthy young adults using structural and diffusion-weighted MRI data. Our findings revealed significant associations between intelligence and network controllers located within the frontal, temporal and parietal cortex. Furthermore, we discovered that these controllers collectively enable access to both easy- and difficult-to-reach network states, aligning with the predictions made by the network neuroscience framework. Additionally, our research demonstrated that the identified network controllers are primarily localized within the left hemisphere and do not reside within regions or connections that possess the highest capacity for structural control in general. This discovery suggests that the identified regions may facilitate specialized control operations and motivates further exploration of the network topology and dynamics underlying intelligence in the human brain.

**Summary:** This study examines the relationship between regional measures of network controllability and general intelligence within a sample of 275 healthy young adults using structural and diffusion-weighted MRI data. We report that individual differences in general intelligence are associated average and modal controllability in specific left-hemisphere cortical regions, and further show that controller regions associated with intelligence are distinct from regions with the highest, centrality, controllability, or communication. These findings reveal a significant structural role for individual regions in controlling the trajectory of the connectome, advancing our understanding of the nature and mechanisms of network controllability in general intelligence.

## 1 Introduction

Research in network neuroscience has advanced our understanding of the network architecture of human intelligence (for a review, see Barbey et al.^1^). The human connectome, comprising a rich constellation of interconnected brain networks, exhibits a highly efficient and specialized organization.^2^ This organization can be characterized as a modular hierarchy, where distinct brain regions form modules that work together while maintaining their individual functional specialization. Within these modules, the connectivity is densely interconnected, enabling rapid information transfer and integration. Simultaneously, long-range connections bridge different modules, facilitating communication and coordination between specialized regions.^3,4^ This network organization enables efficient information flow, supporting complex cognitive processes and facilitating the seamless integration of diverse brain functions.^5–10^

Emerging evidence demonstrates that the system-wide coordination of brain networks is made possible by network flexibility and the capacity to transition between network states, engaging specific brain networks in response to task demands.^11,12^ A central and yet largely unexplored premise of this perspective is the idea that network state transitions can be *controlled* —that brain network topology is organized so that certain regions can more efficiently route information across the cortex^13^ or control the brain’s dynamic reconfiguration in a goal-directed manner.^14^ The relationship between network communication and control in the human connectome is complex,^15–17^ motivating investigations of network control that examine the set of possible network transitions afforded by the brain’s structural organization. Techniques from network control theory allow precise characterization of a brain region’s ability to control transitions between network states—elucidating how individual differences in network topology constrain functional activity and transitions in the service of goal-directed behavior.^18–21^ Network control theory therefore provides a computational framework for measuring and modeling network flexibility,^22^ by applying systems engineering principles^23^ to represent controllability patterns as a set of trajectories between discrete functional states.

The Network Neuroscience Theory of Intelligence^24^ predicts that general intelligence (*g*) depends on the ability to dynamically reorganize brain network topology in a flexible and adaptive manner. Brain network flexibility facilitates performance across diverse cognitive tasks, providing a neurobiological account of how shared variance in performance across tasks may facilitate general intelligence and the positive manifold.^25^ This network understanding of brain structure and function moves beyond previous descriptions focused on localizing cognitive abilities within specific brain regions^26^ or individual brain networks.^27,28^ Instead, this framework proposes that network flexibility is made possible by network controllers that move the system into specific network states, enabling solutions to familiar problems by accessing nearby, easy-to-reach network states and adapting to novel situations by engaging distant, difficult-to-reach network states.^24,29^ Thus, a key claim of the Network Neuroscience Theory is that the global network organization of human intelligence relies on structural brain network controllability and the capacity to access both easy- and difficult-to-reach network states. Previous work has demonstrated individual differences in the controllability of specific network control regions,^14,22,30^ suggesting that structural topology and controllability provide a novel lens for understanding individual differences in intelligence.

The preset work examines structural network controllability in a large sample of healthy young adults (*N* = 275) who underwent structural and diffusion tensor imaging.^31,32^ Subjects completed a comprehensive cognitive battery designed to investigate individual differences in general intelligence.

Network control theory defines three controllability metrics through which a given region may control network transitions across the connectome: average, modal, and boundary controllability. *Average controllability* reflects the overall structural interconnectivity of a region, capturing its ability to steer the connectome into many different (and commonly accessible) state spaces. It assumes equiprobable states, but extensions consider unequal probabilities,^33^ leading to the development of *modal controllability*. Modal controllability identifies regions with the ability to steer the system into less commonly employed, more difficult-to-reach states. Modal controllability has been most studied in neurobiology contexts^34,35^ with several frontoparietal hub regions implicated in general cognitive ability identified as important for transitioning into difficult-to-reach states.^36–39^ In contrast, *boundary controllability* identifies regions with diverse connectivity, reflecting the ability to both integrate and segregate multiple discrete neural communities. While this control operation may play a key role facilitating brain network flexibility,^40^ current theories do not associate boundary controllability with human intelligence, providing a comparison variable against average and modal controllability in our analyses.

Network neuroscience evidence also suggests that intelligence depends on flexible global connectome organization and topology, such that distributed, weakly-connected edges play a key role in facilitating network states required for intelligence.^29,41,42^ This perspective aligns with evidence for a distinction between the role of topological hub regions in facilitating network communication, and the importance of non-hubs for facilitating network control.^15^ The present analysis also aims to determine if regions that control the connectome and produce intelligence are specific; namely, do they overlap with rich-club regions that are highly central to the connectome (supporting communication through strong ties^43^), and do they overlap with regions that exert the highest amount of control across the connectome?

In summary, the present study sought to elucidate the relationship between general intelligence and structural controllability, with a focus on three primary aims: (1) To uncover specific brain regions that drive network control in general intelligence; (2) To establish the specific control operations that are necessary (average, modal, or boundary controllability); and (3) To investigate whether the observed control regions are specific to intelligence, or overlap with regions that are important for communication or control across the connectome in general.

## 2 Methods

### 2.1 Participants

This study was approved by carried out in accordance with the recommendations of the University of Illinois at Urbana-Champaign Institutional Review Board. All subjects gave written informed consent in accordance with the Declaration of Helsinki. Data were collected from a sample of healthy young adults in Illinois as part of a larger study investigating the efficacy of multimodal interventions to enhance fluid intelligence.^32,44,45^ This study reports pre-intervention resting-state fMRI data that were acquired prior to the intervention (and therefore not affected by the larger project). A total of 453 subjects enrolled in this phase of the study, of which approximately two thirds (*N* = 304) were randomly assigned to receive neuroimaging. After removing subjects with incomplete DTI or structural imaging acquisitions, or who attritted from the study between the two days of neuropsychological testing, 275 subjects remained for analysis. All subjects gave written informed consent in accordance with the Declaration of Helsinki. Study inclusion criteria recruited adults (1) aged 18– 44 years; (2) fluent in English; (3) possessing at least a high-school diploma; (4) with normal or corrected to normal vision and hearing; (5) free of any psychoactive medication; (6) without history of neurological, psychological, or endocrine disease; (7) without history of concussion for the past 2 years; (8) not having learning disorders; (9) not smoking > 10 cigarettes a day; (10) with a body mass index < 35; and (11) with at least one positive response on the revised Physical Activity Readiness Questionnaire.^46^ All subjects reported in the present analysis were randomly assigned to brain imaging data collection and were right-handed.

### 2.2 Cognitive Battery

To characterize general intelligence, we analyzed a diverse battery of pre-intervention measures employed in the INSIGHT cognitive assessment protocol.^31,47^ The cognitive tests administered in the battery are presented below.

#### Figure Series

Figure Series is a test of nonverbal reasoning and fluid intelligence abilities similar to the Raven’s Progressive Matrices.^48^ Participants are shown a series of figures with one item missing from the sequence.^49^ The subject’s task is to deduce the rule governing the series and select the option that correctly completes the series. The task is comprised of 30 items in total with a time limit of 60 seconds per item. The primary performance measure for this task is the timed accuracy within a 30-minute limit.

#### LSAT Logical Reasoning

The Logical Reasoning portion of the Law School Admissions Test is used to measure verbal reasoning abilities. Participants are shown word problems posing a logical question. Participants must use reasoning skills and select the best answer of five answer options.^50^ The LSAT is comprised of 25 items in total. The primary outcome measure is the number of items answered correctly within the 35-minute time limit. Previous factor analytic research within the INSIGHT battery supports the test’s association with general intelligence.^44,51^

#### Shipley-2 Vocabulary

The vocabulary subscale of the Shipley Institute of Living Scale is a well-validated measure of crystallized intelligence.^52^ Each item presents a word and asks the participant to select the nearest synonym. The battery consists of 140 items, and is administered with a time limit of 10 minutes. The primary outcome measure for this task is the total number of correct items (i.e., total accuracy).

#### Adult Decision-Making Competence

Decision-making is a critical component of adaptive reasoning and problem-solving. The human judgment and decision-making literature indicates that the ability to resist biases in decision-making relies on three core competencies: (1) comprehension, the capacity to assess the likelihood and value of possible actions and their consequences, (2) integration, the ability to combine available information to make an adaptive choice, and (3) meta-cognitive awareness, the capacity to use analysis and deliberation to evaluate intuitive responses in decision makingbaron2008, kahneman2011, stanovich2010. The A-DMC is designed to assess the consistency and/or accuracy of individuals’ decisions across six subscales: Resistance to Framing, Recognizing Social Norms, Under/Overconfidence, Applying Decision Rules, Risk Perception, and Resistance to Sunk Costs.^47,53^ Prior research further demonstrates that performance on the A-DMC is highly related to general intelligence (with a construct-level correlation of 0.91^54^), suggesting it provides a valid measure for intelligence in the present study.

### 2.3 Estimating the General Factor of Intelligence

The positive manifold (*g* ; general intelligence) emerges from the shared variance across performance on cognitive tasks.^25^ We deploy *exploratory factor analysis* (EFA) to determine the best-fitting number of latent factors for capturing variance among our tasks, employing a bootstrapping approach to validate this solution before finally repeating *confirmatory factor analysis* (CFA) across the complete dataset and generating theoretical factor scores reflecting *g* for each subject. We deploy the bootstrapping procedure train EFA models with oblimin rotation on a random 70% training set of participant data, and then test that model against the 30% holdout to investigate relative goodness-of-fit of a 1- and 2-factor solution through maximum likelihood estimation, repeating this procedure 10,000 times for each model. This approach allows us to predict individual differences in our latent measures of intelligence while accounting for individual measurement errors in manifest variables.

### 2.4 Brain Imaging

Diffusion Tensor Imaging (DTI) assesses water diffusion in the brain, allowing characterization of white matter integrity and simulation of structural tractography. As part of a comprehensive two-hour battery of multimodal neuroimaging acquisition, DTI data was collected with a Siemens 3 Tesla scanner using a 32-channel head coil. 30-directional DWI data were acquired on the half shell at b = 1000 *s/mm*^2^. Data were acquired at 21.91.9*mm* across 72 slices using single-shot, spin-echo EPI acquisition with an echo time (TE) of 100 ms, a repetition time (TR) of five seconds, and SMS multiband and GRAPPA at 2.^55–57^ Structural T1-weighted MPRAGE^58^ was also acquired at 0.9 mm isotropic with TE = 2.32 ms and TR = 1.9 seconds.

Tractography characterizes the strength of long-range white matter projections between pairs of brain regions. DTI processing began by processing BIDS-formatted MRI data using qsiprep^59^ to extract subject-space tissue maps from structural and DTI acquisitions. Structural preprocessing and parcellation was performed through FSL^60^ and Freesurfer.^61^ Cortical parcellation was performed through Freesurfer’s recon-all stream^61,62^ to derive volumetric cortical and subcortical labels in FSL’s Desikan anatomical space.^63^ Resulting regional masks were registered into diffusion space (referencing fractional anisotropy (FA) images) using bbregister initialized with FLIRT.^64–66^ FSL’s topup and eddy were used to correct DTI images for inhomogeneity in the scan acquisition while accounting for potential subject motion.^67–70^ DTI tractography was performed using FSL’s diffusion tensor stream, using dtifit-derived mean FA images as registration targets when registering structural masks and confound signals (white matter; WM, cerebrospinal fluid; CSF) to DTI image space. GPU-accelerated bedpostx was used to fit voxelwise tensors reflecting diffusion gradient after bvecs rotation..^71,72^ FSL’s probtrackx2^73,74^ performed GPU-accelerated probabilistic tractography by seeding 5000 streamlines from each voxel within a mask for each anatomical region of interest (ROI). The analysis retained all streamlines successfully propagated to an incident ROI, with an exclusion mask employed to discount all streamline that tracked into CSF.

In this way, probabilistic tractography characterizes the structural topology of white-matter connections. For each subject, tractography outputs square matrix (82 × 82) of absolute streamline connectivity between each gray matter ROI. To normalize for stochastic differences in the total number of connected streamlines derived from differences in the number of source and target voxels, we averaged raw streamline connectivity for each edge by the total volume of both incidents’ ROIs. This procedure produced a weighted and undirected structural connectome that estimates the density of white matter connectivity between each pair of regions (for further details, see Sharp et al.^45^). To normalize for bidirectional sampling of undirected edges between each incident region, each resulting final connectome was averaged across the diagonal.

### 2.5 Structural Network Controllability Analysis

Network control theory is a mathematical framework for modeling network flexibility and the capacity of a system to traverse between network states. Accumulating evidence indicates that network control measures predict individual differences in brain network re-configuration^75^ when applied to the structural topology of specific white matter connections.^30^ The present work employs an existing toolbox for structural network controllability to characterize average, modal, and boundary controllability of each structural brain region in subjects’ connectomes.^22^

Formally, a network can be represented as the graph 𝒢= (𝒱, ℰ) where 𝒱 represents a set of nodes and ℰ a set of existing directed edges. If N represents the number of nodes, the symmetric weighted adjacency matrix **A** = [*a*_*ij*_], **A** ∈ ℝ^*N* ×*N*^ would define the network topology, where *a*_*ij*_ denotes the weights of the acyclic edges in ℰ (i.e., *A*_*ii*_ = 0 ∀*i*). Then the following discrete time-invariant network model can be used to derive the state at the next timepoint:

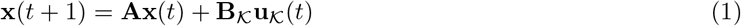

where **x** : ℕ_≥0_ → ℝ^*N* ×*N*^ denotes the time-dependent state of brain regions and **A** denotes the adjacency matrix. 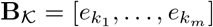 denotes the input matrix (where *e*_*i*_ is the *i*-th canonical vector and |*e*_*i*_| = *N*) associated with the control nodes 𝒦 := {*k*_1_, …, *k*_*m*_} ⊂ 𝒱, while **u**_𝒦_ : ℕ_≥0_ → ℝ^*m*^ denotes the control signal injected into the network via the nodes 𝒦. Then, the controllability matrix is defined as:

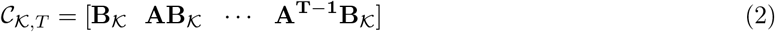

The network 𝒢 is controllable in *T* steps by the control nodes 𝒦 *if and only if* 𝒞_𝒦,*t*_ is full row rank. Alternatively, a network 𝒢 is controllable in *T* steps by the control nodes 𝒦 *if and only if* the *controllability Grammian* 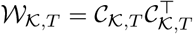 is positive definite. We used the following notions of controllability as in Gu et0020al.:^22^

*Average Controllability* is the average energy from the input/control nodes, representing the ability of a node to control each mode of the dynamics of a network. We use Trace(𝒲_𝒦,*T*_) instead of Trace 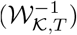 for computational convenience and and because the two measures are strongly correlated.

*Modal Controllability* is the ability of a node to control each mode of the dynamics of a network. Let *λ*_1_(**A**), …, *λ*_*N*_ (**A**) denote the modes from the control node *i*. Then, we define:

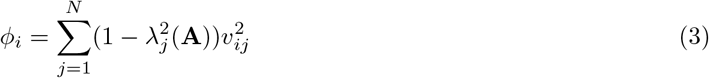

the scaled measure of the controllability of the modes in the region *i*, where **V** = [*v*_*ij*_] is the eigenvector matrix of **A**. A value of *v*_*ij*_ = 0 implies that the mode *λ*_*j*_(**A**) is not controllable from *i*.

*Boundary Controllability* measures the capacity of a node to connect disparate brain regions. We assign a boundary controllability score of 1 or 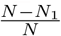 to the set of *N*_1_ boundary nodes determined by the robust partition of the network. We complement this set with the additional boundary nodes determined by the partitions of the least controllable subnetwork, as determined by the Fiedler eigenvector of the graph 𝒢 Laplacian.

Additionally, we generate an average sample-level structural connectome by averaging connectivity across all subjects and distance-transforming the resulting edge strengths. Based on this connectome, sample-level node and edge betweenness centrality were used to identify structural rich-club nodes and the highest-connectivity structural edges. Structural region centrality was contrasted with region controllability at the sample level. At the subject level, edgewise lesioning of the top 10% highly connected sample-level edges was used to recalculate subject’s regional controllability metrics in a simulation analysis, to identify regions where network control was most dependent on high network communication.

To relate regional network controllability to individual differences in latent measures of cognitive ability, we perform linear regression to predict subjects’ general intelligence based on each region’s network controllability properties (i.e., three univariate analyses) after controlling for age, sex, ESL (English as a second language) status, and years of education. We corrected all p-values from these regressions for multiple comparisons by keeping the false discovery rate (FDR; step-up BH following Benjamini and Hochberg^76^ at *α* = 0.05).

## 3 Results

### 3.1 Cognitive Battery

The descriptive statistics and correlations between the administered tests were computed to examine the relationship between each of the cognitive measures. Table 1 presents the mean, median, and standard deviation for each of the administered tests. Consistent with prior reports,^29,44,51^ we observed that performance on each subtest of the A-DMC was higher in this sample than the validation sample studied by Bruine de Bruin et al.^47^ The logical reasoning portion of the LSAT displayed the highest pearson’s correlation with A-DMC battery subscores, (range: *r* = [0.12 − 0.48]), followed by the vocabulary test, (range: *r* = [0.05 − 0.40]), with the Figure Series measure showing the weakest associations (range: *r* = [0.03 − 0.37]).

**Table 1:** Descriptive statistics for cognitive assessment battery

| Variable | $\mu$ | $\tilde{x}$ | $\sigma$ |
| --- | --- | --- | --- |
| Figure Series | 2.05 | 1.98 | 1.50 |
| LSAT | 12.6 | 13 | 4.06 |
| Shipley 2 Vocab | 0.47 | 0.51 | 0.22 |
| Applying Decision Rules | 0.82 | 0.87 | 0.16 |
| Resistance to Sunk Costs | 4.31 | 4.30 | 0.70 |
| Consistency in Risk Perception | 0.74 | 0.75 | 0.74 |
| Resistance to Framing | 4.18 | 4.21 | 4.29 |
| Recognizing Social Norms | 0.47 | 0.51 | 0.22 |
| Over/Underconfidence | 1.07 | 1.07 | 0.12 |

Bartlett’s test of sphericity confirms the variance of our cognitive battery is amenable to factor analysis (df=28, *χ*^2^ = 376, *p <* .001), as does the Kaiser, Meyer, Olkin measure (*KMO* = 0.79). Multiple factor estimation measures equivalently supported a 1- or 2-factor solution as optimal, while none supported a 3-factor solution. Horn’s parallel analysis was in favor of a 1-factor solution. We contrasted exploratory factor analyses identifying both a 1-factor and a 2-factor solution for modeling latent variance in our cognitive battery. Under a 70/30 training/test split, bootstrapped *p* − *values* from *χ*^2^ tests of model fit for the 1- and 2-factor solutions were similar for each model (10,000 folds, average *p* = .12 and *p* = .14 respectively).

Furthermore, *χ*^2^ tests comparing between the nested 1- and 2-factor EFA models were reliably nonsignificant (*p* = .32). These results suggest that both EFA solutions performed adequately for modeling the factor structure of our holdout cognitive testing data under bootstrapping, and that no significant difference (across many *χ*^2^ tests) existed between the resulting bootstrapped CFA models. Collectively, these results suggest that a 1-factor solution is well-supported and most parsimonious for our data. In keeping with the generally accepted psychometric structure of intelligence,^25^ we labeled the variable in the 1-factor solution *g*, reflecting general cognitive ability.

Based on this analysis, we elected to proceed with SEM modeling of a 1-factor solution (Table 2). We performed CFA for a 1-factor solutions in our sample via maximum likelihood estimation implemented in R’s lavaan package.^77^ Fit indices were high (1-factor: *χ*^2^ = 51.3, *p* = 0.003, *RMSEA* = 0.055, *CFI* = 0.936, *r*^2^ = 0.24).

**Table 2:**
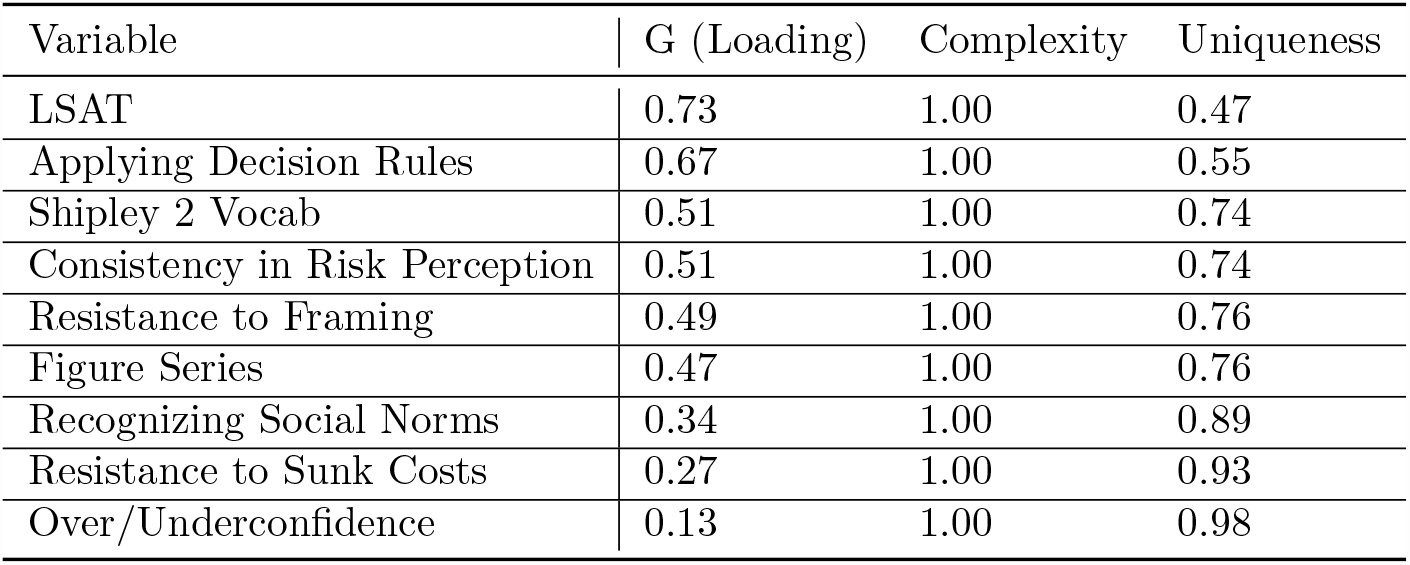
1-factor solution loadings for general intelligence (g)

### 3.2 Structural Controllability of *g*

Structural network controllability was calculated for each region in the Desikan Atlas using normalized streamline density estimates from white matter tractography. Each streamline was normalized with respect to the total volume of its two incident ROIs, and each normalized structural connectome was then averaged across the diagonal to produce an undirected matrix accounting for stochastic differences in directional streamline propagation. Average, modal, and boundary controllability were calculated for each ROI in each subject, then averaged for a sample-wide connectome across all individuals. We regressed individual differences in structural network controllability against estimates of intelligence by fitting regression models for each region and factor score and correcting for multiple comparisons via FDR (*N* = 275, FDR *α* = 0.05). After FDR-correction, individual differences in general intelligence (*g*) are reliably associated with the average and modal controllability of multiple predominantly left-hemisphere structural ROIs (Table 3). The average structural controllability of six regions is positively and significantly associated with g, in brain areas distributed across temporal and parietal cortices. Average controllability in these regions reflects the ability to transition the brain into many equiprobable states. Average controllability of these regions individually account for between 4-7% of the variance in general intelligence. Modal controllability of the left inferior parietal and left cingulate isthmus also show a significant correlation with general intelligence, reflecting the ability to transition the connectome into difficult to reach network configurations. FDR-correction identified these regressions under a critical *p*− *values* of 0.004. No boundary controllability regions survived FDR-correction.

**Table 3:** Network controllability of general intelligence (*g*)

| Region | Controllability | $r^2$ | $p$ | Centrality | Controllability Rank |
| --- | --- | --- | --- | --- | --- |
| Left Caudal Anterior Cingulate | Average | 0.06 | <0.0001 | 47/82 | 66/82 |
| Left Inferior Parietal | Average | 0.05 | <0.002 | 36/82 | 67/82 |
| Left Inferior Temporal | Average | 0.05 | <0.002 | 29/82 | 3/82 |
| Left Isthmus Cingulate | Average | 0.05 | <0.002 | 68/82 | 77/82 |
| Left Middle Temporal | Average | 0.04 | <0.003 | 45/82 | 26/82 |
| Right Inferior Temporal | Average | 0.06 | <0.0002 | 17/82 | 60/82 |
| Left Inferior Parietal | Modal | 0.07 | <0.0001 | 36/82 | 53/82 |
| Left Isthmus Cingulate | Modal | 0.05 | <0.006 | 68/82 | 81/82 |

Notably, none of the regions where controllability is significantly associated with intelligence are members of the structural rich-club at the sample level (i.e., none appear in the top 10% of regions ranked by centrality; Table 3). This dissociation highlights that regions capable of facilitating cognition via control operations are not the ones most strongly implicated in global communication, suggesting that network control operations important to intelligence are originating in regions with specialized connectivity profiles. Further, seven out of eight significant regions do not appear in the top 20% of regions with the largest controllability statistic (Table 3), suggesting that control operations in these regions enable network states and transitions that are specific to intelligence, instead of reflecting greater control over the connectome in general. These results suggest that general intelligence relies on specialized network controllers that move the system into specific network states, enabling solutions to familiar problems by accessing nearby, easy-to-reach network states (average controllability) and adapting to novel situations by engaging distant, difficult-to-reach network states (modal controllability).

To further assess the relationship between average controllability and network communication, we lesioned the highest communication edges (top 10% betweenness centrality) in each subject’s connectome and calculated resulting decreases to structural controllability. This analysis examines whether average controllability engages strongly-vs weakly connected edges and whether regions of high centrality are implicated (i.e., that afford high communication in a rich-club structural organization).

Table 4 demonstrates that lesioning edgewise communication across the connectome produces varying decreases to structural controllability for regions associated with *g* (Δ = [1% − 65%]; *μ* = 28%). In contrast, the highest controllability regions (Table 5) show more substantial decreases to communication (Δ = [55% − 98%]; *μ* = 84%). Contrasting these findings suggests that average structural controllability associated with *g* does not primarily depend on strongly connected, high-communication edges, suggesting that individual differences in average control operations may instead reflect weak edges or more fine-scale aspects of connectome organization. These findings also illustrate that regional average controllability values associated with *g* may be one to two orders of magnitude lower than the largest controllability observed across the connectome, again suggesting that controllability operations associated with *g* reflect profiles of structural organization that may be specialized for general intelligence.

**Table 4:** Change in Lesioned Average Controllability (General Intelligence Regions)

| Region | Delta | % change | Centrality Rank |
| --- | --- | --- | --- |
| Left Caudal Anterior Cingulate | -0.44 | -29.7% | 47 |
| Left Inferior Parietal | -0.25 | -37.2% | 36 |
| Left Inferior Temporal | 0.04 | -1.0% | 29 |
| Left Isthmus Cingulate | -1.61 | -30.0% | 68 |
| Left Middle Temporal | 0.22 | -64.7% | 45 |
| Right Inferior Temporal | 0.06 | -3.5% | 17 |

**Table 5:** Change in Lesioned Average Controllability (Highest Centrality Regions)

| Region | Delta | % change | Centrality Rank |
| --- | --- | --- | --- |
| Left Pericalcarine | -49.9 | -63.7% | 1 |
| Left Parahippocampal | -258.3 | -93.6% | 2 |
| Left Amygdala | -197.3 | -67.0% | 3 |
| Right Hippocampus | -39.6 | -85.8% | 4 |
| Right Accumbens | -208.7 | -54.9% | 5 |
| Right Cuneus | -57.9 | -94.2% | 6 |
| Left Temporal Pole | -69.8 | -97.5% | 7 |
| Right Thalamus | -206.4 | -97.2% | 8 |
| Left Transverse Temporal | -146.8 | -97.9% | 9 |
| Right Caudal Anterior Cingulate | -185.6 | -92.8% | 10 |

## 4 Discussion

Network neuroscience theory predicts that intelligence arises from system-wide network dynamics and the capacity to flexibly transition between network states. According to this view, network flexibility is made possible by network controllers that move the system into specific network states, enabling solutions to familiar problems by accessing nearby, easy-to-reach network states and adapting to novel situations by engaging distant, difficult-to-reach network states. Although this framework predicts that general intelligence depends on network controllability, the specific cortical regions that serve as network controllers and the nature of their control operations remain to be established. To investigate these issues, the present study examined the relationship between regional measures of network controllability and general intelligence within a sample of 275 healthy young adults using structural and diffusion-weighted MRI data. We report several main findings that advance our understanding of the nature and mechanisms of network controllability in general intelligence.

### Network Control Regions

Our findings revealed significant associations between intelligence and network controllers located within the frontal, temporal, and parietal cortex. Previous research has associated executive control and intelligence with the organization of these regions in the functional^38,78,79^ and structural^80–82^ domain, findings replicated by our localization of controller regions for general intelligence. Previous work has also shown that controllability in some of these regions is associated with variance on individual cognitive tasks, including a task of fluid ability.^83^ Our report extends this finding and is the first to demonstrate that individual differences in controllability are associated with general intelligence overall, highlighting a potential network mechanism through which the positive manifold may emerge.

Additionally, our findings revealed that the identified network controllers are primarily localized within the left hemisphere. Prior large-scale lesion studies of human intelligence have observed a distributed network of left-lateralized brain regions,^84^ consistent with the observed pattern of findings. Evidence of functional asymmetry in human and animal cortex is well-established,^85^ with left-hemisphere function broadly implicated in language and reasoning. Hemispheric asymmetries in cognitive control have been demonstrated through lesion studies, such that unilateral right-hemisphere lesion is associated with control deficits to attention and external processing,^86,87^ whereas left-hemisphere lesion is associated with control deficits in verbal and non-verbal cognitive performance.^79,88^ Our results extend these findings by showing that left-hemisphere structural organization may be specialized to support multiple structural control operations implicated in human intelligence, either through their neuroanatomical or topological configuration.

Further, we demonstrate that control operations do not reside within regions or connections that possess the highest capacity for structural control in general. This discovery suggests that the identified regions may facilitate specialized control operations and motivates further exploration of the network topology and dynamics underlying intelligence in the human brain.

### Network Control Operations

Our findings revealed that structural controllers collectively enable access to both easy- and difficult-to-reach network states, aligning with the predictions made by the network neuroscience framework. We show that general intelligence is associated with modal and average controllability and therefore enables diverse transitions into both easy- and difficult-to-reach network states, consistent with the predictions of the Network Neuroscience Theory.^24^ Previous research has associated high average controllability with performance on individual cognitive tasks.^83^ Our findings extend this work by demonstrating a role for both average and modal controllability in facilitating cognitive performance. These findings suggest that network flexibility is made possible by network controllers transitioning the connectome into different network states, either enabling solutions to familiar problems by accessing nearby, easy-to-reach network states^5,9^ or adapting to novel situations by engaging distant, difficult-to-reach network states.^14,34,89^

### Network Control Theory: Future Directions

Research demonstrates that human intelligence is shaped by the structural brain connectome, facilitating globally coordinated and dynamic connectivity across brain networks. Our findings reveal a significant structural role for individual regions in controlling the trajectory of the connectome to support human intelligence.

Here, we report that individual differences in general intelligence are associated with specific network control operations in specific cortical regions. Our results show that general intelligence is significantly related to individual differences in the average or modal controllability of multiple predominantly left-hemisphere cortical regions; and that controller regions associated with intelligence are a distinct set from regions with the highest, centrality, controllability, or communication across the connectome overall.

Given the relative paucity of empirical results demonstrating relationships between structural controllability and individual differences in cognition, further validation studies should be undertaken to substantiate these findings in additional populations. A potential limitation of our findings is the relatively small portion of variance in intelligence explained by individual regional measures of structural controllability. Our approach here affords specificity, localization, and control of type I error. These findings do however motivate a more comprehensive multivariate or predictive modeling investigation into structural controllability, for example, at the level of mesoscale brain networks.

The discovery of predominant left-hemisphere lateralization of control regions suggests they may be anatomically specialized to facilitate control operations and dynamics, a claim supported by the dissociation between controller regions for general intelligence and highly-connected or central regions in general. This finding motivates further exploration of the network topology and dynamics underlying intelligence, in particular to characterize how multivariate or community-level structural topology shapes functional dynamics and constrains transitions between network states. Where do control nodes for general intelligence reside within the hierarchical organization of the connectome? How do they interact with within-community or between-community structure to enable local or global network transitions? Are control regions the causal source of network transitions, or are they the topological consequence of a well-organized connectome?^90,91^ The relationship between controllability and the positive manifold motivates future multimodal imaging studies that leverage the relationship between structure and function, for example joint structure-function modeling that considers dynamics in the context of underlying structural connectivity networks. Addressing these questions promises to shed light on mechanisms that allow the brain to arrive at precise network configurations and produce the breadth of human cognitive ability.

## Acknowledgement

We are grateful to the INSIGHT investigators and project team, especially our project manager Patricia Jones, as well as the numerous fellows, students, and staff that made the INSIGHT project possible. This research is based upon work supported by the Office of the Director of National Intelligence (ODNI), Intelligence Advanced Research Projects Activity (IARPA), via Contract 2014–13121700004 to the University of Illinois Urbana-Champaign (PI: Barbey) as well as the National Science Foundation Grant IIS-2123781. The views and conclusions contained herein are those of the authors and should not be interpreted as necessarily representing the official policies or endorsements, either expressed or implied, of the ODNI, IARPA, or the U.S. Government. The U.S. Government is authorized to reproduce and distribute reprints for Governmental purposes notwithstanding any copyright annotation thereon.

## Data & Code Availability

Restrictions made by study participants during informed consent on the distribution or future research uses of their deidentified data prevent open distribution of the dataset used in this paper. Data to reproduce analyses contained in this paper are available from the authors on request. Code to perform tractography, factor analysis, and controllabiltiy analysis is available at https://github.com/eandrsn2/struct-control-g.

## Author Contributions

Conceptualization, Methodology: E.D.A., L.R.V., B.H., P.D.R., A.N.K., R.R.W., C.E.Z., B.K., and A.K.B.; Software, Investigation, Data Curation, E.D.A.; Writing—Original Draft: E.D.A., L.R.V, P.D.R.; Writing—Review & Editing: E.D.A., L.R.V., B.H., and A.K.B.; Resources, Project Administration, and Funding Acquisition: A.K.B., L.R.V.; Supervision, L.R.V., B.K., and A.K.B.

## Declaration of Interests

The authors declare no competing interests.

## Notes

### Competing Interest Statement

The authors have declared no competing interest.

